# Multivariate adaptive shrinkage improves cross-population transcriptome prediction for transcriptome-wide association studies in underrepresented populations

**DOI:** 10.1101/2023.02.09.527747

**Authors:** Daniel S. Araujo, Chris Nguyen, Xiaowei Hu, Anna V. Mikhaylova, Chris Gignoux, Kristin Ardlie, Kent D. Taylor, Peter Durda, Yongmei Liu, George Papanicolaou, Michael H. Cho, Stephen S. Rich, Jerome I. Rotter, NHLBI TOPMed Consortium, Hae Kyung Im, Ani Manichaikul, Heather E. Wheeler

## Abstract

Transcriptome prediction models built with data from European-descent individuals are less accurate when applied to different populations because of differences in linkage disequilibrium patterns and allele frequencies. We hypothesized methods that leverage shared regulatory effects across different conditions, in this case, across different populations may improve cross-population transcriptome prediction. To test this hypothesis, we made transcriptome prediction models for use in transcriptome-wide association studies (TWAS) using different methods (Elastic Net, Joint-Tissue Imputation (JTI), Matrix eQTL, Multivariate Adaptive Shrinkage in R (MASHR), and Transcriptome-Integrated Genetic Association Resource (TIGAR)) and tested their out-of-sample transcriptome prediction accuracy in population-matched and cross-population scenarios. Additionally, to evaluate model applicability in TWAS, we integrated publicly available multi-ethnic genome-wide association study (GWAS) summary statistics from the Population Architecture using Genomics and Epidemiology Study (PAGE) and Pan-UK Biobank with our developed transcriptome prediction models. In regard to transcriptome prediction accuracy, MASHR models performed better or the same as other methods in both population-matched and cross-population transcriptome predictions. Furthermore, in multi-ethnic TWAS, MASHR models yielded more discoveries that replicate in both PAGE and PanUKBB across all methods analyzed, including loci previously mapped in GWAS and new loci previously not found in GWAS. Overall, our study demonstrates the importance of using methods that benefit from different populations’ effect size estimates in order to improve TWAS for multi-ethnic or underrepresented populations.

## 1. INTRODUCTION

Through genome-wide association studies (GWAS), many associations between single nucleotide polymorphisms (SNPs) and diverse phenotypes have been uncovered^1^. However, most GWAS to date have been conducted on individuals of European descent, even though they make up less than one fifth of the total global population^2, 3^. Ancestry diversity in human genetic studies is important because as linkage disequilibrium and allele frequencies differ among populations, associations found within European ancestry individuals may not reflect associations for individuals of other ancestries and vice versa^3^. Some efforts to increase ancestry diversity in human genetics studies include the NHLBI Trans-Omics for Precision Medicine (TOPMed) consortium^4^, the Population Architecture using Genomics and Epidemiology (PAGE) study^5^, the Human Heredity and Health in Africa (H3Africa) initiative^6^, and the Pan-ancestry genetic analysis of the UK Biobank (PanUKBB^7^).

Alongside GWAS, transcriptome-wide association studies (TWAS) test predicted gene expression levels for association with complex traits of interest, identifying gene-trait associated pairs^8^. Different TWAS methods, such as PrediXcan and FUSION, work by estimating gene expression through genotype data using transcriptomic prediction models built on expression quantitative trait loci (eQTL) data^9,10^. Similarly to GWAS, TWAS are also negatively affected by ancestry underrepresentation, as gene expression prediction models for use in TWAS are often trained in European descent datasets, which reduces the power of studies conducted with individuals of other ancestries^11^,^12^. Still, we expect the underlying biological mechanisms of complex traits to be shared across human populations^11^,^13^, and thus prediction methods that account for allelic heterogeneity and better estimate effect sizes can improve the discovery rate and interpretation of TWAS across populations.

Here, we used genomic and transcriptomic data from the Multi-Ethnic Study of Atherosclerosis (MESA)^14^ multi-omics pilot study of TOPMed to build TWAS prediction models (Figure 1). Using five different methods to estimate effect sizes, Elastic-Net^15^,^16^, Joint-Tissue Imputation (JTI)^17^, Matrix eQTL^18^, multivariate adaptive shrinkage (MASHR)^19^, and Transcriptome-Integrated Genetic Association Resource (TIGAR)^20^, we built population-specific transcriptomic prediction models for four MESA-defined populations – African American, Chinese, European, and Hispanic/Latino – across three blood cell types and evaluated their prediction performance in the Geuvadis^21^ cohort using PrediXcan^9^. From there, we used S-PrediXcan^22^ to apply our models to GWAS summary statistics of complex traits from the multi-ethnic PAGE^5^ study and PanUKBB^7^. We hypothesized that MASHR and JTI were most likely to improve transcriptome prediction and increase the number of TWAS hits in comparison to the other methods, as they both leverage similar effect size estimates across different conditions - in this case, different populations - to adjust effect sizes. In agreement to that, our results indicated that in cross-population predictions, MASHR models have a higher transcriptome prediction accuracy than other methods. Furthermore, in our TWAS, MASHR models discovered the highest number of associated gene-trait pairs across all population models. These findings illustrate that leveraging genetic diversity and effect size estimates across populations can help improve current transcriptome prediction models, which may increase discovery and replication in association studies in underrepresented populations or multi-ethnic cohorts.

**Figure 1:**
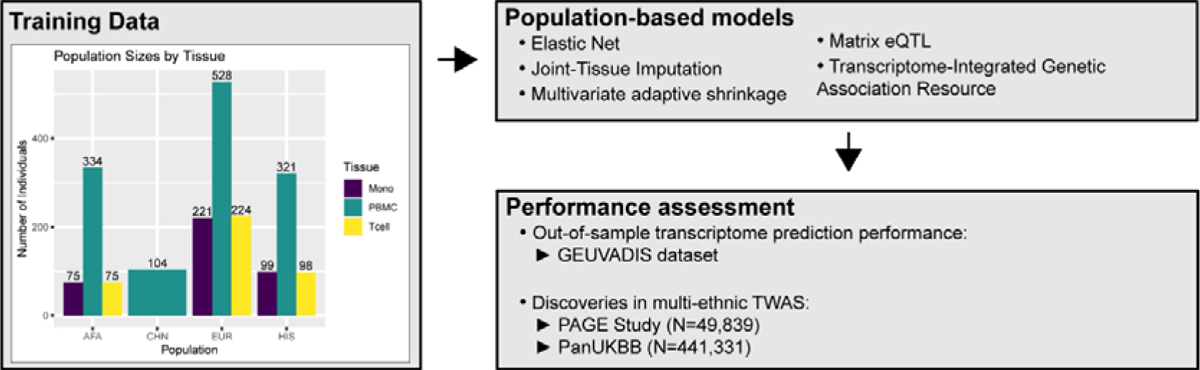
Overall study methodology. Using TOPMed MESA as a training dataset, we built population-based transcriptome prediction models using five different methods (Elastic Net (EN), Joint-Tissue Imputation (JTI), Multivariate adaptive shrinkage (MASHR), Matrix eQTL, and Transcriptome-Integrated Genetic Association Resource (TIGAR)). With these transcriptome models, we evaluated their out-of-sample transcriptome prediction accuracy using the GEUVADIS dataset. Additionally, we assessed their applicability in multi-ethnic TWAS using GWAS summary statistics from the PAGE Study and PanUKBB. AFA = African American, CHN = Chinese, EUR = European, HIS = Hispanic/Latino.

## 2. METHODS

### a. Training dataset

To build our transcriptome prediction models, we used data from the Multi-Ethnic Study of Atherosclerosis (MESA)^14^ multi-omics pilot study of the NHLBI Trans-Omics for Precision Medicine (TOPMed) consortium. This data set includes genotypes derived from whole genome sequencing and transcripts per million (TPM) values derived from RNA-Seq for individuals of four different populations – African American (AFA), Chinese (CHN), European (EUR), and Hispanic/Latino (HIS) – for three different blood cell types: peripheral blood mononuclear cells (PBMC, ALL n = 1287, AFA n = 334, CHN n = 104, EUR n = 528, HIS n= 321), CD16+monocytes (Mono, ALL n = 395, AFA n = 75, EUR n = 221, HIS n = 99), and CD4+ T-cells (T cells, ALL n = 397, AFA n = 75, EUR n = 224, HIS n = 98).

### b. Genotype and RNA-Seq QC

We performed QC on each MESA tissue-population pair separately. For the genotype data^4^ (Freeze 8, phs001416.v2.p1), we excluded INDELs, multi-allelic SNPs, and ambiguous-strand SNPs (A/T, T/A, C/G, G/C), and removed the remaining variants with MAF < 0.01 and HWE < 1 x 10^-6^ using PLINK^23^ v1.9. For chromosome X, filtering by HWE was only applied in variants found within the pseudoautosomal regions based on GRCh38 positions. Furthermore, for the non-pseudoautosomal region of X, male dosages were assigned either 0 or 2. After QC, the average numbers of non-ambiguous SNPs remaining per population across all cell types were: AFA = 15.7M; CHN = 8.4M; EUR = 9.7M; HIS = 13.2M.

For the RNA-Seq data, we also performed QC separately by tissue-population. First, we removed genes with average TPM values < 0.1. For some individuals, RNA expression levels were measured at two different time points (Exam 1 and Exam 5); thus, after log-transforming each measurement and adjusting for age and sex as covariates using linear regression and extracting the residuals, we took the mean of the two time points (or the single adjusted log-transformed value, if expression levels were only measured once), performed rank-based inverse normal transformation, and adjusted for the first 10 genotype and 10 expression PCs. To estimate principal components, we used PC-AiR^24^ with kinship threshold of ∼0.022, which corresponds to 4^th^ degree relatives. No individuals were removed. For each tissue, we removed genes absent in at least one population. After QC, we had 17,585 genes in PBMC, 14,503 in Mono, and 16,647 in T cells. We used GENCODE^25^ annotation v38 to annotate gene types (e.g. protein-coding, lncRNA, etc.) and gene transcription start and end sites.

### c. Gene expression cis-heritability estimation

We estimated gene expression heritability (h^2^) using cis-SNPs within the 1Mb region upstream of the transcription start site and 1Mb region downstream of the transcription end site. Using the genotype data filtered only by HWE P-value < 1 x 10^-6^, for each tissue-population pair, we first performed LD-pruning with a 500 variants count window, a 50 variants count step, and a 0.2 r^2^ threshold using PLINK^23^ v1.9. Then, for each gene, we extracted cis-SNPs and excluded SNPs with MAF < 0.01. Finally, to assess cis-SNP expression heritability, we estimated the genetic relationship matrix and h^2^ using GCTA-GREML^26^ with the “--reml-no-constrain” option. We considered a gene heritable if it had a positive h^2^ estimate (h^2^ - 2*S.E. > 0.01 and p-value < 0.05) in at least one MESA population. In total, 9,206 genes were heritable in PBMC, 3,804 in Mono, and 4,053 in T cells. We only built transcriptome prediction models for these heritable genes across all populations in their respective cell types.

### d. Transcriptome prediction models

With the aforementioned genotype and gene expression data, we built transcriptome prediction models for each MESA tissue-population pair, and for each gene we considered cis-SNPs as defined in the previous section. Additionally, we only considered SNPs present in the GWAS summary statistics of the Population Architecture using Genomics and Epidemiology (PAGE) study^5^ to build our prediction models to make sure that there would be a high overlap between SNPs in the transcriptome models and SNPs in the GWAS summary statistics. After merging with PAGE SNPs, the average numbers of SNPs left in our dataset were: AFA = 12.8M; CHN = 6.2M; EUR = 7.4M; HIS = 10.5M.

We built our population-based models using five different approaches. The first was elastic-net (EN) regression using the *glmnet* package in R^15, 16^, with mixing parameter α = 0.5. We considered EN as our baseline model, as it has been previously used to make transcriptome prediction models for the TOPMed MESA data^27^.

The second method implemented was mash (Multivariate Adaptive Shrinkage)^19^ in R (MASHR). Unlike EN, MASHR does not estimate weights by itself; rather, it takes z-score (or weight and standard error) matrices as input and adjusts them based on correlation patterns present in the data in an empirical Bayes algorithm, allowing for both shared and condition-specific effects. By doing so, MASHR increases power and effect size estimation accuracy^19^. Originally, MASHR applicability was demonstrated by leveraging effect size estimates across different tissues^19^, however, herein we sought to assess its potential to leverage effect sizes across populations. We ran MASHR for each gene at a time, using cis-SNPs weights (effect sizes) estimated by Matrix eQTL^18^ and MESA populations as different conditions (Figure 2A). Then, we split MASHR-adjusted weights according to their respective populations, and selected the top SNP (lowest local false sign rate) per gene to determine which SNPs would end up in the final models (Figure 2B). Local false sign rate is similar to false discovery rate, but it is more rigorous as it also takes into account the direction of effect^19^. Thus, by selecting one top SNP per population, the maximum number of SNPs per gene in the final model is 4, which corresponds to the number of populations in our study. If two or more populations had the same variant as top SNP, it was only included once. To make population-based models, we used population-specific effect sizes, taken from the corresponding MASHR output matrices.

**Figure 2:**
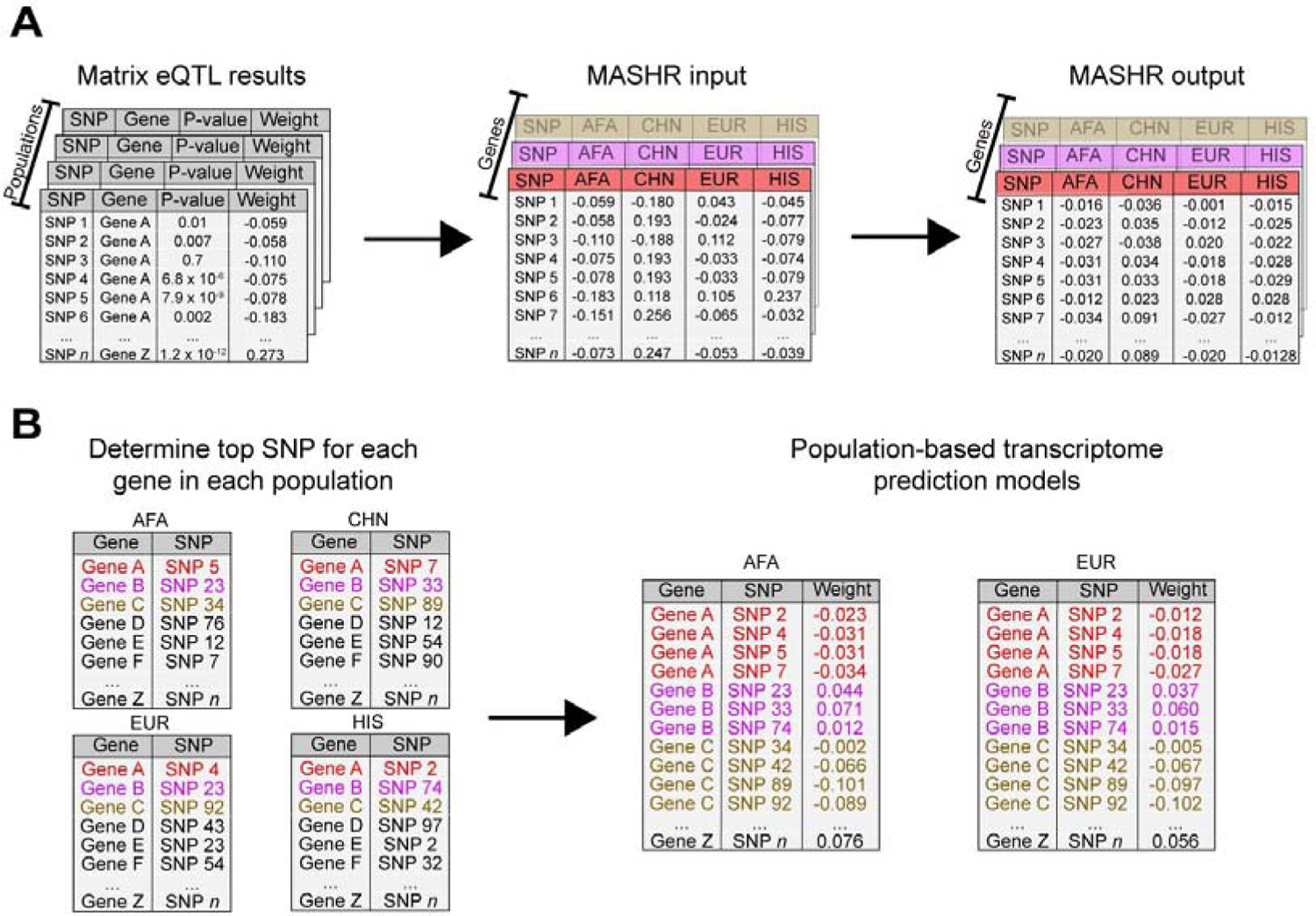
Design of the methodology implemented to make MASHR models. (A) Using effect sizes estimated using Matrix eQTL within each population dataset, we combined them across genes, with the different populations as conditions, to use as input for MASHR. The output matrixes contain adjusted effect sizes. (B) For each population, we selected the top SNP (lowest local false sign rate) per gene. Then, we concatenated the Gene-top SNP pairs across populations to determine which SNPs would end up in the final models. Lastly, to make our population-based transcriptome prediction models, we used population-specific effect sizes, taken from the corresponding MASHR output matrices. AFA = African American, CHN = Chinese, EUR = European, HIS = Hispanic/Latino.

The third method was based on the unadjusted effect sizes estimated by Matrix eQTL^18^ using the linear regression model. We used the same approach taken to build the MASHR models, including the SNP with the lowest p-value from each population, but the key difference is that we made the models using the unadjusted effect sizes.

The fourth method we used was Transcriptome-Integrated Genetic Association Resource (TIGAR), which trains transcriptome imputation models using either EN or nonparametric Bayesian Dirichlet Process Regression (DPR)^20^. As we already used EN to make a set of transcriptome prediction models, we opted to make DPR-based models. We used TIGAR’s default parameters to train our models, such as using the Variational Bayesian algorithm and outputting fixed effect sizes. However, by default, TIGAR performs 5-fold cross validation (CV) during training, and only outputs results if the final average CV R² is equal or greater than 0.005; thus, since we did not implement CV for any of the aforementioned methods and instead tested performance in an independent sample, we opted to skip this step of TIGAR’s pipeline and generate outputs for all genes. Most gene models generated by TIGAR had hundreds of SNPs with near-zero effect sizes. To reduce memory requirements for storage of these models, we removed SNPs with effect sizes smaller than 1 x 10^-4^.

The fifth and last method we implemented was Joint-Tissue Imputation (JTI)^17^. JTI was designed to leverage similarity in gene expression and DNase 1 hypersensitive sites across different tissues to possibly improve prediction performance. Thus, similarly to MASHR, we sought to assess whether the method could be adapted to use populations instead of tissues. To assess gene expression similarity between MESA populations, we computed transcriptome-wide pairwise correlations between populations using the median TPM value per gene. Additionally, we did not have population DNase 1 hypersensitivity site data, so we set column five to 1 in our input files. By default, JTI performs 5-fold CV and only produces outputs for genes with average CV R greater than 0.1. Thus, similarly to TIGAR, we removed this filtering step of the pipeline to generate output for all genes regardless of CV performance.

To perform TWAS using GWAS summary statistics data, it is necessary to have information about the correlation between the SNPs used to predict gene expression levels^22^. Thus, for all our transcriptome prediction models previously mentioned, we computed pairwise covariances for the SNPs within each TOPMed MESA population model using the respective population dosage data. All model files are freely available for anyone to use (see Data Availability section).

### e. Assessing transcriptome prediction performance

To evaluate the gene expression prediction performance of all our transcriptome prediction models, we used DNA and lymphoblastoid cell lines RNA-Seq data from 449 individuals in the Geuvadis^21^ study. Individuals within the testing dataset belong to five different populations (Utah residents with Northern and Western European ancestry (CEU), n = 91; Finnish in Finland (FIN), n = 92; British in England and Scotland (GBR), n = 86; Toscani in Italy (TSI), n = 91; Yoruba in Ibadan, Nigeria (YRI), n = 89), which we analyzed both separately and together (ALL). Similarly to our training dataset, we performed rank-based inverse normal transformation on the gene expression levels, and adjusted for the first 10 genotype and 10 expression PCs, using the residuals as observed expression levels. With the Geuvadis genotype data and our transcriptome prediction models, we used PrediXcan^9^ to estimate gene expression levels. PrediXcan is a two-step TWAS method, in which the first step is to estimate genetically regulated expression levels (GReX). Thus, to assess transcriptome prediction performance, we compared GReX to the adjusted, measured expression levels using Spearman correlation.

### f. Assessing performance in transcriptome-wide association studies

To test the applicability of our transcriptome prediction models in multi-ethnic association studies, we applied S-PrediXcan^22^ to GWAS summary statistics from the Population Architecture using Genomics and Epidemiology (PAGE) study^5^. The PAGE study consists of 28 different phenotypes tested for association with variants within a multi-ethnic, non-European cohort of 49,839 individuals (Hispanic/Latino (n=22,216), African American (n=17,299), Asian (n=4,680), Native Hawaiian (n=3,940), Native American (n=652) or Other (n=1,052)). Since we tested multiple phenotypes and transcriptome prediction models in our TWAS, we used a conservative approach and considered genes as significantly associated with a phenotype if the association p-value was less than the standard Bonferroni corrected GWAS significance threshold of 5 x 10^-8^.

To replicate the associations found in PAGE, we also applied S-PrediXcan^19^ to PanUKBB^7^ GWAS summary statistics (N=441,331; European (n=420,531), Central/South Asian (n=8,876), African (n=6,636), East Asian (n=2,709), Middle Eastern (n=1,599) or Admixed American (n=980)). For similarity purposes, we selected summary statistics of phenotypes that overlap with the ones tested in PAGE (Table S1). As previously described, a gene-trait pair association was considered significant if its p-value was less than the Bonferroni corrected GWAS significance threshold of 5 x 10^-8^. Furthermore, we deemed significant gene-trait pair associations as replicated if they were detected by the same MESA tissue-population model and had the same direction of effect in PAGE and PanUKBB. To assess if the gene-trait association pairs reported in our study are novel or not, we compared them to studies found in the GWAS Catalog^1^ (All associations v1.0.2 file downloaded on 11/9/2022).

## 3. RESULTS

### a. Increased sample sizes improve gene expression cis-heritability estimation

With the goal of improving transcriptome prediction in diverse populations, we first determined which gene expression traits were heritable and thus amenable to genetic prediction, using genome-wide genotype and RNA-Seq data from three blood cell types (PBMCs, monocytes, T cells) in TOPMed MESA. We estimated cis-heritability (h²) using data from four different populations (African American - AFA, Chinese - CHN, European - EUR, and Hispanic/Latino - HIS). Variation in h² estimation between populations is expected due to differences in allele frequencies and LD patterns; however, we show that larger population sample sizes yield more significant (p-value < 0.05) h² estimates (Figure 3). Using the PBMC dataset as an example, with the EUR dataset (n = 528), we assessed h² for 10,228 genes, however, we estimated h² for 8,765 genes using the AFA dataset (n = 334) (Figure 3A).

**Figure 3:**
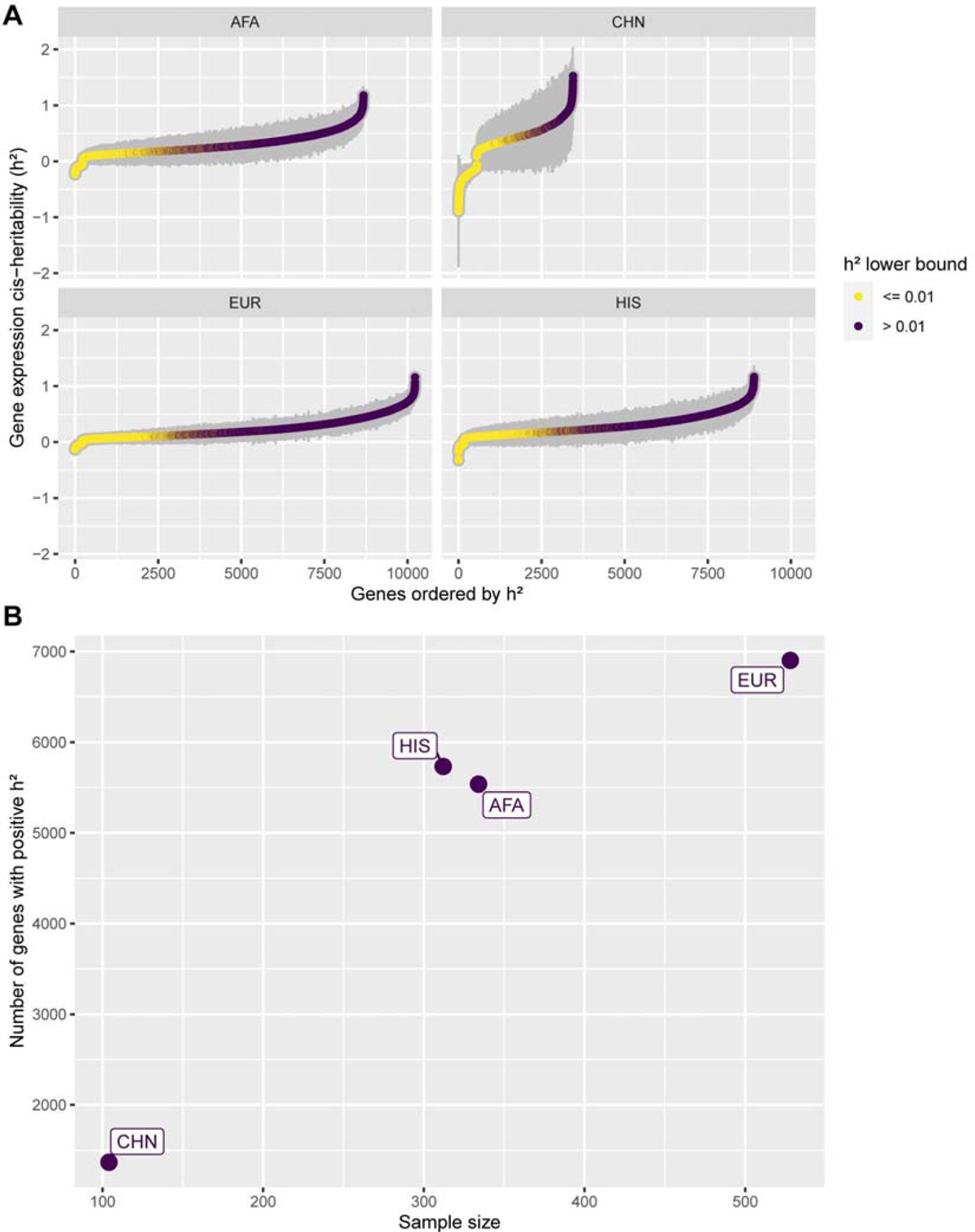
PBMC gene expression cis-heritability estimates across MESA populations. (A) Gene expression cis-heritability (h²) estimated for different genes across different MESA population datasets in PBMC. Only genes with significant estimated h² (p-value < 0.05) are shown. Gray bars represent the standard errors (2*S.E.). Genes are ordered on the x-axis in ascending h² order, and colored according to the h² lower bound (h² - 2*S.E.). (B) Number of significant heritable genes (p-value < 0.05 and h² lower bound > 0.01) within each PBMC population dataset, by sample size. AFA = African American, CHN = Chinese, EUR = European, HIS = Hispanic/Latino.

Moreover, we see a great impact on the CHN population, which has the smallest sample size. For that population, we managed to estimate h² for only 3,448 genes. The same pattern repeats when analyzing only the heritable genes (h² lower bound > 0.01). In EUR, 6,902 genes were deemed heritable, whereas in AFA and CHN the amount of heritable genes is 5,537 and 1,367, respectively (Figure 3B). Thus, larger sample sizes are needed to better pinpoint h² estimates, especially in non-European populations. In total, analyzing the union across all populations’ results, we detected 9,206 heritable genes in PBMCs, 3,804 in monocytes, and 4,053 in T Cells.

### b. MASHR models improve cross-population transcriptome prediction performance

To improve TWAS power for discovery and replication across all populations, we sought to improve cross-population transcriptome prediction accuracy. For this, we used data from four different populations and built gene expression prediction models using five different methods (Elastic Net (EN), Transcriptome-Integrated Genetic Association Resource (TIGAR), Matrix eQTL, multivariate adaptive shrinkage in R (MASHR), and Joint-Tissue Imputation (JTI)). We chose EN as a baseline approach for comparison in our analysis, as it has been previously shown to have better performance than other common machine learning methods such as random forest, K-nearest neighbor, and support vector regression^28^. Furthermore, we trained gene expression prediction models by applying TIGAR’s nonparametric Bayesian Dirichlet Process Regression pipeline^20^. Using Matrix eQTL, we estimated univariate effect sizes for each cis-SNP-gene relationship and we developed an algorithm to include top SNPs from each population, but population-estimated effect sizes in each population’s model (Figure 2). Matrix eQTL effect sizes are the input for MASHR, which we hypothesized might better estimate cross-population effect sizes, due to its flexibility in allowing both shared and population-specific effects^19, 29^. Similarly, JTI was designed to leverage correlation across different tissues to improve gene expression prediction^17^; thus, we also adapted its pipeline to perform cross-population leveraging. By filtering our models to include only genes with positive h² (h² lower bound > 0.01) in at least one population, we saw that among all methods used, we obtained more gene models in MatrixeQTL and MASHR (Figure 4A). The difference is especially greater in the CHN population model.

**Figure 4:**
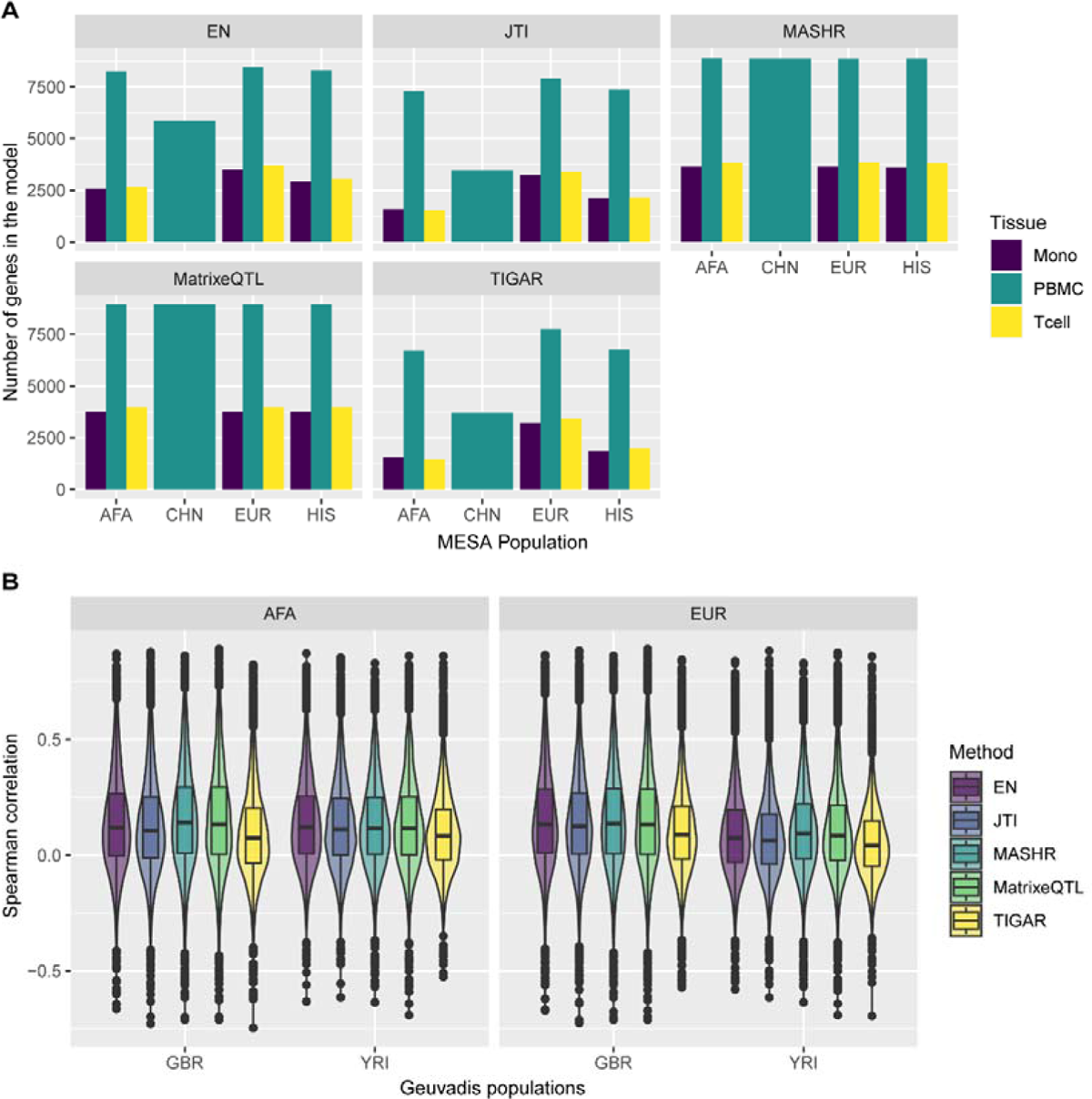
Comparison of MESA population transcriptome prediction models. (A) The number of genes in each MESA population model, by method and tissue. (B) Prediction performance (Spearman’s rho) of EN, JTI, MASHR, MatrixeQTL, and TIGAR PBMC MESA population models in Geuvadis GBR and YRI populations. Only the intersection of genes with expression predicted by all methods for each MESA-Geuvadis population pair are shown. MASHR performed better than or the same as all other methods (see Table S2 for all pairwise comparisons).

To evaluate model performance at population-matched and cross-population transcriptome predictions, we used data from the Geuvadis study, which comprises individuals of West African or European descent. We defined “population-matched predictions” as the scenarios in which the transcriptome model MESA training data and Geuvadis test data have the closest genetic distance with available data, and we defined “cross-population predictions” as any other pairs (Figure S1). Overall, across all Geuvadis populations, the methods tested show distinct performances (Figure S2). This result, however, may be influenced by the fact that different transcriptome models have a different number of genes in them (Figure 4A). Thus, we sought to compare performances considering the intersection of genes with expression predicted by all methods. Focusing on Geuvadis GBR and YRI populations, which have similar sample sizes and are of distinct continental ancestries, we observed that MASHR models significantly outperform the other methods in cross-population transcriptome predictions, as seen in the AFA-GBR and EUR-YRI MESA-Geuvadis populations pairs (Figure 4B, Table S2). The only exception is in AFA-GBR, in which MASHR and MatrixeQTL have similar performances. Additionally, in population-matched scenarios (AFA-YRI and EUR-GBR), prediction performance does not significantly differ between MASHR, MatrixeQTL, and EN. All three aforementioned methods significantly outperform JTI and TIGAR in population-matched predictions (Table S2). Moreover, we also performed pairwise comparisons between all methods using all Geuvadis populations, taking into account the intersection of genes with expression predicted in each case. Overall, across all MESA transcriptome models and Geuvadis populations, MASHR models either performed better or the same as other methods in both population-matched or cross-population transcriptome prediction scenarios (Table S3).

### c. Leveraging effect sizes across different populations improves discovery rate in multi-ethnic TWAS

In order to investigate the applicability of the models we built in multi-ethnic TWAS, we used S-PrediXcan with GWAS summary statistics of complex traits from PAGE and PanUKBB. We show that across all tissue-population models, MASHR identified the highest number of gene-trait pair associations (208) that replicated in both PAGE and PanUKBB (P < 5 x 10^-8^), followed by Matrix eQTL (173), JTI (131), EN (94), and TIGAR (91) (Table S3). When analyzing the total number of discoveries separately for each population, MASHR had the highest number of gene-trait pairs in most population models (Figure 5A). The only exception is with HIS models, in which both MASHR and MatrixeQTL had the same number of discoveries. The discovery rate improvement by MASHR is exceptionally high in CHN models, as it had almost twice the number of discoveries as the second-highest method (27 by MASHR vs. 14 by MatrixeQTL).

**Figure 5:**
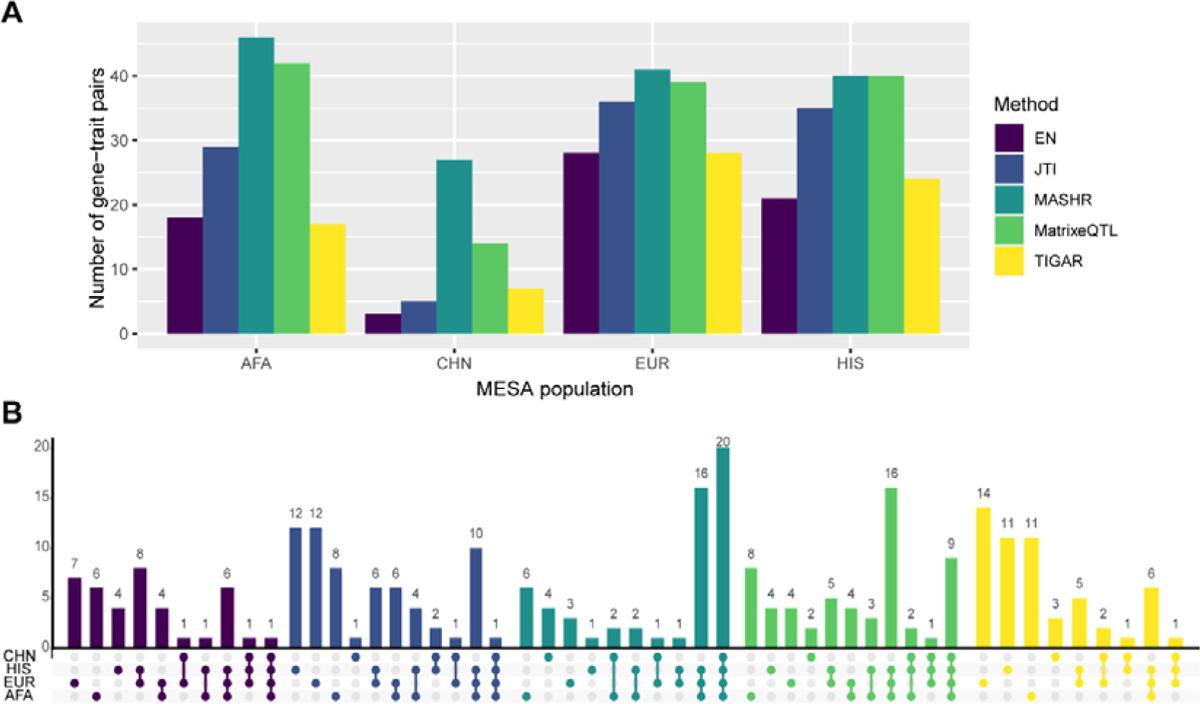
Number of significant S-PrediXcan gene-trait pairs in PAGE and PanUKBB GWAS summary statistics. (A) Total number of significant gene-trait pairs discovered by each MESA population model (considering the union of the three tissues), by method. (B) Number of significant gene-trait pairs discovered with individual or multiple MESA populations colored by method (considering the union of the three tissues). Population set intersections are indicated on the x-axis in color.

Additionally, when comparing gene-trait pairs, we saw that most MASHR hits were shared between population models, whereas other methods have higher population-specific discoveries (Figure 5B). Most MatrixeQTL hits were also shared by many population models, but not to the same degree as MASHR. Altogether, these findings indicate that MASHR models show high consistency and also suggest that TWAS results are not as affected by the MASHR population model used as compared to other methods.

To contextualize our models’ findings, we investigated whether the discovered gene-trait pairs had been previously reported in any studies in the GWAS Catalog (https://www.ebi.ac.uk/gwas/home). We saw that across 105 distinct gene-trait pairs associations found (totaling 697 across all models), 38 (36.19%) have not been reported in the GWAS Catalog, and therefore may be novel associations that require further investigation (Table S4). Out of those potential new biological associations, most of them (13) were discovered with MASHR AFA models (Table S4). Furthermore, out of the 67 distinct known GWAS catalog associations discovered, MASHR models identified most of them (Table S3). For instance, MASHR EUR models found 34 known associations, followed by MASHR AFA with 33, and MatrixeQTL EUR with 32 (Figure S4).

## 4. DISCUSSION

In this work, we sought to build population-based transcriptome prediction models for TWAS using data from the TOPMed MESA cohort using five distinct approaches. We saw that although the AFA and HIS populations’ datasets contained the highest numbers of SNPs after quality control, EUR yielded the highest number of gene expression traits with significant heritability estimates across all tissues analyzed. This is most likely due to the higher sample size in EUR in comparison to AFA and HIS, as larger sample sizes provide higher statistical power to detect eQTLs with smaller effects^30^. Furthermore, we saw that the number of genes in each population transcriptome model is not the same across all methods tested. Some transcriptome prediction models, such as the ones built using EN or JTI, only contain genes for which the SNPs effect sizes converged during training, which is not a limiting factor for MASHR, MatrixeQTL, and TIGAR. One of the factors that impacts the number of genes for which SNPs effect sizes converge during training is sample size, which explains the lower number of genes in the EN and JTI CHN model in comparison to other population models.

Furthermore, although sample size does not impact the number of gene models trained for TIGAR in the same degree as EN and JTI, it influences SNP effect size estimation^31^. Thus, when we removed SNPs with near-zero effects, there was a drop in the number of genes in the final population transcriptome models for TIGAR. Test data sample size has also been shown to positively correlate with gene expression prediction accuracy^32^.

In addition to sample size, gene expression prediction accuracy is known to be greater when the training and testing datasets have similar ancestries^11, 12, 32, 33^; however, non-European ancestries are vastly underrepresented in human genetics studies^2, 3^, which compromises the ability to build accurate TWAS models for them. Thus, using data from the Geuvadis cohort, we evaluated the transcriptome prediction performance of our models and found that MASHR models either significantly outperformed all other methods tested, or had similar performance. Previous studies have shown that by borrowing information across different conditions, such as tissues^19^ or cell types^34^, MASHR identifies shared- or condition-specific eQTLs, which can enhance causal gene identification^29^, as well as improve effect size estimation accuracy^19^.

Similarly, by leveraging effect size estimates across multiple populations, MASHR improved cross-population transcriptome prediction without compromising population-matched prediction accuracy. Interestingly, another method we tested, JTI, was also originally designed to leverage similarity in gene expression and DNase 1 hypersensitive sites across tissues in order to improve transcriptome prediction accuracy^17^. However, our results showed that it performed worse than MASHR and the same as EN in cross-population transcriptome prediction. This suggests that distinct cross-condition leveraging frameworks may have different performances when applied across populations. One possible reason for differences in performance is that JTI uses EN weighted by condition similarity to estimate effect sizes and select SNPs to be included in the final models, whereas for MASHR, our pipeline selects one SNP per condition. Since more SNPs with less significant effect sizes were included in our EN and JTI models, greater uncertainty in effect sizes likely led to lower transcriptome prediction accuracy compared to MASHR. Furthermore, among the methods evaluated, TIGAR had the lowest prediction performance. Originally, TIGAR was benchmarked against EN, and showed better transcriptome prediction accuracy; however, unlike in our analysis, their analysis included only genes whose expression heritability was equal or lower than 0.2^20^.

Discovery and replication of TWAS associations are also related to the ancestries of the transcriptome prediction model training dataset and ancestries of the TWAS sample dataset^11^. Thus, we assessed the applicability of our models in TWAS using S-PrediXcan on PAGE and PanUKBB GWAS summary statistics and found that across all tissues and populations, MASHR models yielded the highest number of total gene-trait pairs associations, with MASHR AFA reporting the highest number. In this manner, it seems that although MASHR improved gene expression prediction accuracy for all populations analyzed, using transcriptome prediction models that match the ancestries of the GWAS dataset still yields the highest number of TWAS discoveries, which is in agreement with many previous studies^11, 35–38^. Our results also showed that although JTI transcriptome prediction was not as accurate as baseline EN, JTI models had more TWAS discoveries than EN. This exemplifies how integrating data from different genetic ancestries may improve TWAS.

By investigating which associations had been previously reported in the GWAS Catalog, we saw that most new discoveries were found by MASHR models. Some of these possible new discoveries are unique to MASHR models and have been corroborated previously, such as *YJEFN3* (also known as *AIBP2*) and triglycerides, whose low expression in zebrafish increases cellular unesterified cholesterol levels^39^, consistent with our S-PrediXcan effect size directions (PAGE effect size = −0.52, p-value = 6.1 x 10^-16^; PanUKBB effect size = −0.86, p-value = 7.1 x 10^-86^). Additionally, we also saw that MASHR models showed higher consistency across the different population transcriptome prediction models, which means that TWAS results are not as affected by the population model used as other methods.

One limitation of our TWAS is that we used transcriptome prediction models trained in PBMCs, monocytes and T cells, and those tissues might not be the most appropriate for some phenotypes in PAGE or PanUKBB. Additionally, because of the smaller sample sizes for some populations in our training dataset, h² and eQTL effect sizes estimates have large standard errors, which may affect the ability of MASHR to adjust effect sizes across different conditions based on correlation patterns present in the data. Regardless of that, our results mainly demonstrate that we can implement cross-population effect size leveraging using a method first applied to do cross-tissue effect size leveraging - and improve cross-population transcriptome prediction accuracy in doing so. Thus, increasing sample size for underrepresented populations will improve current MASHR TWAS models’ performances, as well as increase genetic diversity in the data. Another TWAS method, METRO, which implements a likelihood-based inference framework to incorporate transcriptome prediction models built on datasets of two different genetic ancestries, has also shown enhanced TWAS power^40^. METRO jointly models gene expression and the phenotype of interest^40^, and thus was not directly comparable to the five methods we tested here, which all separate the transcriptome prediction step from the association test. Given that this traditional two-stage TWAS procedure ignores uncertainty in the expression prediction, the joint approach of METRO across more than two populations is an area of future TWAS methods research. Furthermore, while our study focused on transcriptome prediction, MASHR could also be adapted to possibly improve cross-population polygenic risk scores (PRS). Indeed, other methods like PRS-CSx jointly model complex traits effects across populations in order to improve PRS^41^. MASHR is most useful when population effects are shared, as demonstrated by the more consistent S-PrediXcan results, but population-specific effects are also relevant. For instance, a study in a large African American and Latino cohort discovered eQTLs only present at appreciable allele frequencies in African ancestry populations^38^. Moreover, since our MASHR models focus on the top SNPs, we might not be including enough eQTLs in the models, especially for those genes whose expression is genetically regulated by multiple eQTLs with small effects.

In conclusion, our results demonstrate the importance and the benefits of increasing ancestry diversity in the field of human genetics, especially regarding association studies. As shown, sample size is valuable for assessing gene expression heritability and for accurately estimating eQTL effect sizes, and thus some populations are negatively affected due to the lack of data. However, by making transcriptome prediction models that leverage effect size estimates across different populations using multivariate adaptive shrinkage, we were able to increase gene expression prediction performance for scenarios in which the training data and test data have distant (“cross-population”) genetic distances with available data. Additionally, when applied to multi-ethnic TWAS, the aforementioned models yielded more discoveries across all methods analyzed, even detecting well-known associations that were not detected by other methods. Thus, in order to further improve TWAS in multi-ethnic or underrepresented populations and possibly reduce health care disparities, it is necessary to use methods that consider shared- and population-specific effect sizes, as well as increase available data of underrepresented populations.

## ACKNOWLEDGEMENTS

This work is supported by the NIH National Human Genome Research Institute Academic Research Enhancement Award R15 HG009569 (HEW). Whole genome sequencing (WGS) for the Trans-Omics in Precision Medicine (TOPMed) program was supported by the National Heart, Lung and Blood Institute (NHLBI). WGS for “NHLBI TOPMed: Multi-Ethnic Study of Atherosclerosis (MESA)” (phs001416.v1.p1) was performed at the Broad Institute of MIT and Harvard (3U54HG003067-13S1). Centralized read mapping and genotype calling, along with variant quality metrics and filtering were provided by the TOPMed Informatics Research Center (3R01HL-117626-02S1). Phenotype harmonization, data management, sample-identity QC, and general study coordination, were provided by the TOPMed Data Coordinating Center (3R01HL-120393-02S1), and TOPMed MESA Multi-Omics (HHSN2682015000031/HSN26800004). The MESA projects are conducted and supported by the National Heart, Lung, and Blood Institute (NHLBI) in collaboration with MESA investigators. Support for the Multi-Ethnic Study of Atherosclerosis (MESA) projects are conducted and supported by the National Heart, Lung, and Blood Institute (NHLBI) in collaboration with MESA investigators. Support for MESA is provided by contracts 75N92020D00001, HHSN268201500003I, N01-HC-95159, 75N92020D00005, N01-HC-95160, 75N92020D00002, N01-HC-95161, 75N92020D00003, N01-HC-95162, 75N92020D00006, N01-HC-95163, 75N92020D00004, N01-HC-95164, 75N92020D00007, N01-HC-95165, N01-HC-95166, N01-HC-95167, N01-HC-95168, N01-HC-95169, UL1-TR-000040, UL1-TR-001079, UL1-TR-001420, UL1TR001881, DK063491, and R01HL105756. The MESA Epigenomics and Transcriptomics Studies were funded by National Institutes of Health grants 1R01HL101250, 1RF1AG054474, R01HL126477, R01DK101921, and R01HL135009. The authors thank the other investigators, the staff, and the participants of the MESA study for their valuable contributions. A full list of participating MESA investigators and institutes can be found at http://www.mesa-nhlbi.org.

## 5. DATA AVAILABILITY

All scripts used for analyses, including a pipeline to derive new MASHR models, are available at https://github.com/danielsarj/TOPMed_MESA_crosspop_portability. MESA populations prediction models and raw S-PrediXcan TWAS output files are available at https://doi.org/10.5281/zenodo.7551844. TOPMed MESA data are under controlled access in dbGaP at https://www.ncbi.nlm.nih.gov/gap/ through study accession phs001416.v2.p1. Geuvadis expression data is at Array Express (E-GEUV-1) and genotype data is at http://www.internationalgenome.org/. PAGE GWAS summary statistics are available in the GWAS Catalog at https://www.ebi.ac.uk/gwas/publications/31217584. PanUKBB GWAS summary statistics are available at https://pan.ukbb.broadinstitute.org/phenotypes/index.html.

## Supplementary Data

**Figure S1:**
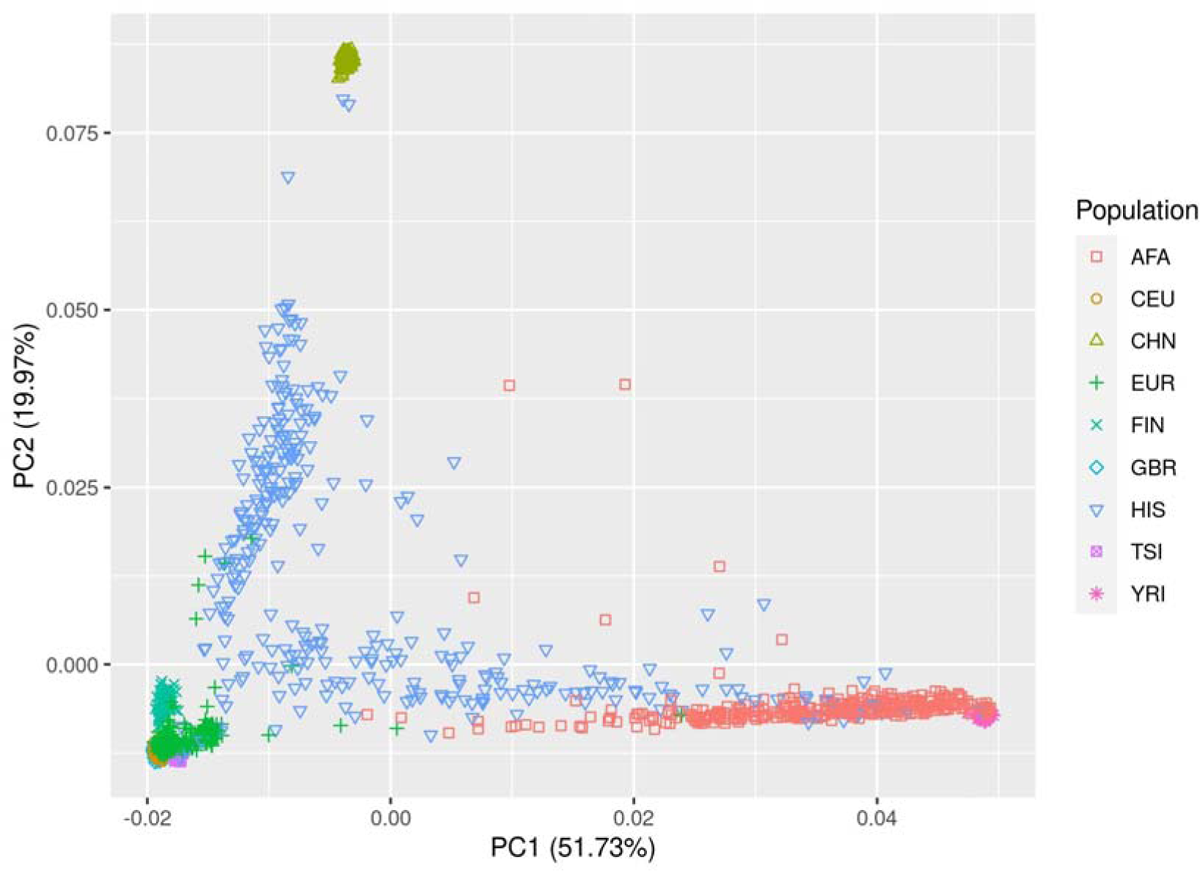
Genotype principal component analysis. Plot of the first two principal components of TOPMed MESA populations with Geuvadis populations. AFA = African American (TOPMed), CEU = Utah residents with Northern and Western European ancestry (Geuvadis), CHN = Chinese (TOPMed), EUR = European (TOPMed), FIN = Finnish in Finland (Geuvadis), GBR = British in England and Scotland (Geuvadis), HIS = Hispanic/Latino (TOPMed), TSI = Toscani in Italy (Geuvadis), YRI = Yoruba in Ibadan, Nigeria (Geuvadis).

**Figure S2:**
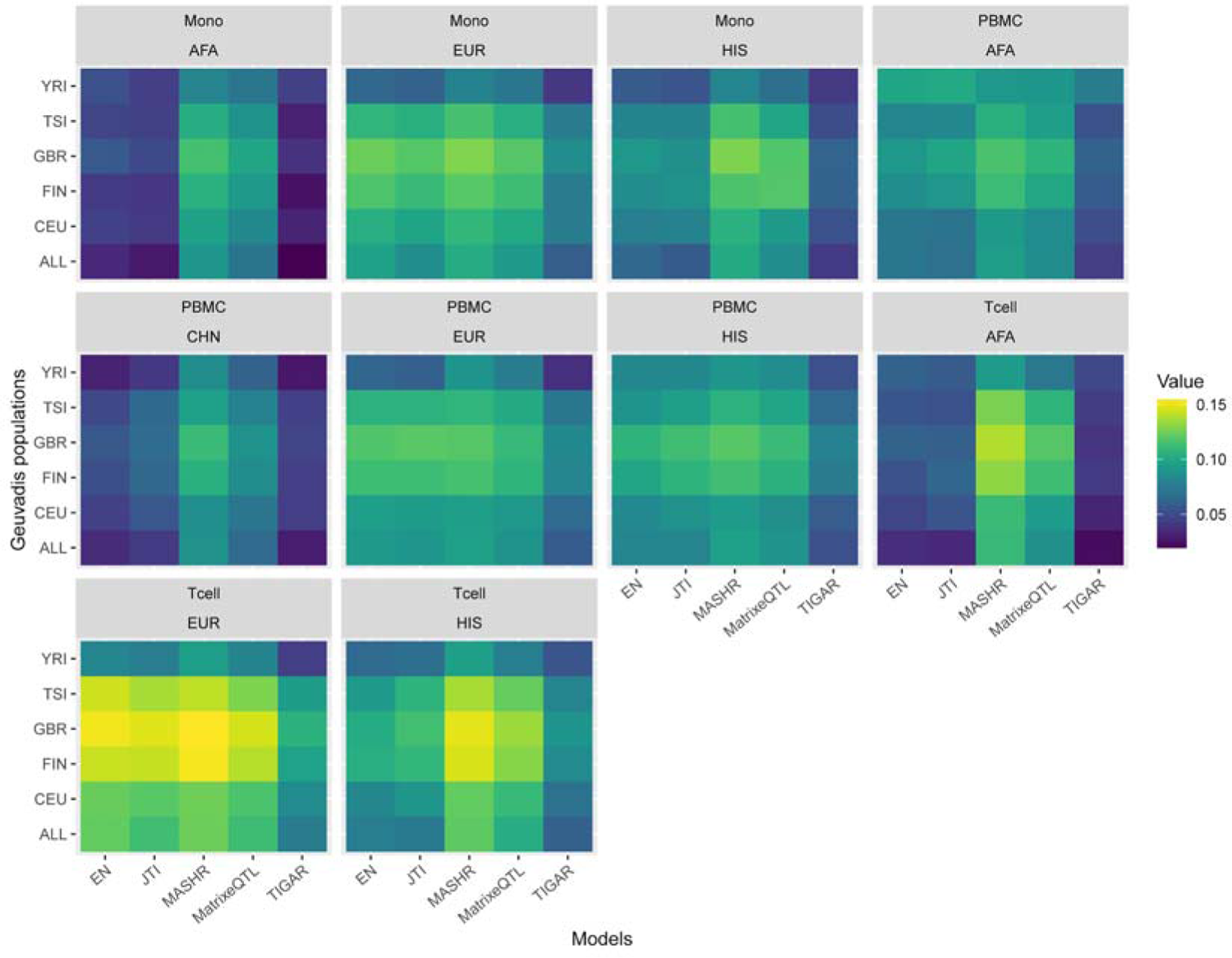
Overall prediction performance of MESA population models in Geuvadis. Prediction performance (median Spearman’s rho) of EN, JTI, MASHR, MatrixeQTL, and TIGAR MESA population models in all Geuvadis populations.

**Figure S3:**
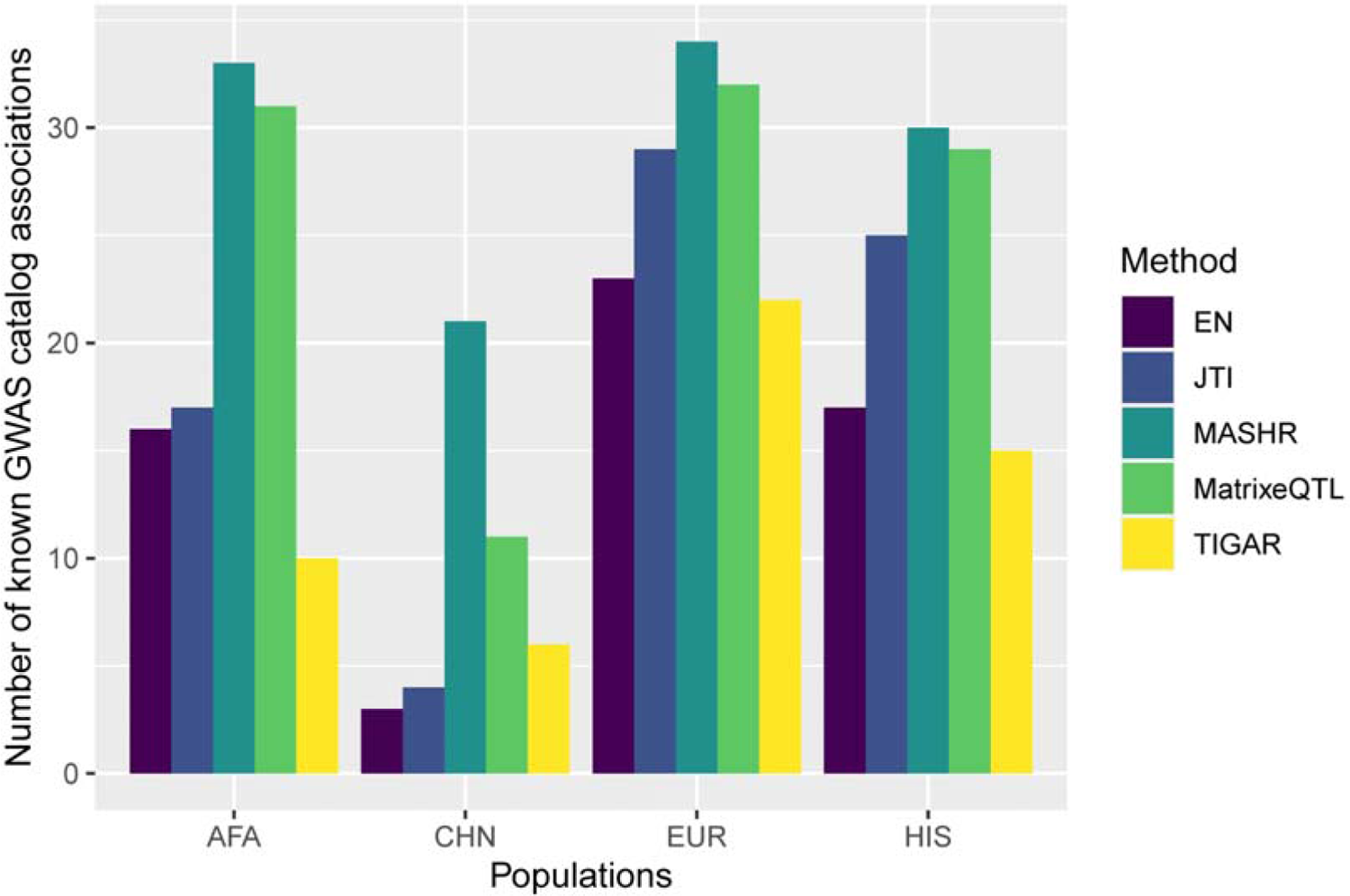
Number of significant S-PrediXcan gene-trait pairs in PAGE and PanUKBB GWAS summary statistics that have been reported in the GWAS catalog. Total number of significant gene-trait pairs discovered by each MESA population model (considering the union of the three tissues), by method.

---

**Table S1:** PAGE and PanUKBB summary statistics used in this study.

**Table S2:** Performance comparisons of PBMC AFA and EUR MESA transcriptome prediction models in the GBR and YRI Geuvadis populations between all methods.

**Table S3:** Pairwise comparisons of the performance of EN, JTI, MASHR, MatrixeQTL, and TIGAR MESA transcriptome prediction models in all Geuvadis populations.

**Table S4:** Compiled S-PrediXcan gene-trait pair discoveries, significant in PAGE and PanUKBB GWAS summary statistics with the same direction of effect.

**Table S5:** List of NHLBI TOPMed consortium members.

