## Supplementary figures and images for "Multivariate adaptive shrinkage improves cross-population transcriptome prediction for transcriptome-wide association studies in underrepresented populations"

### Supplemental Figure 1

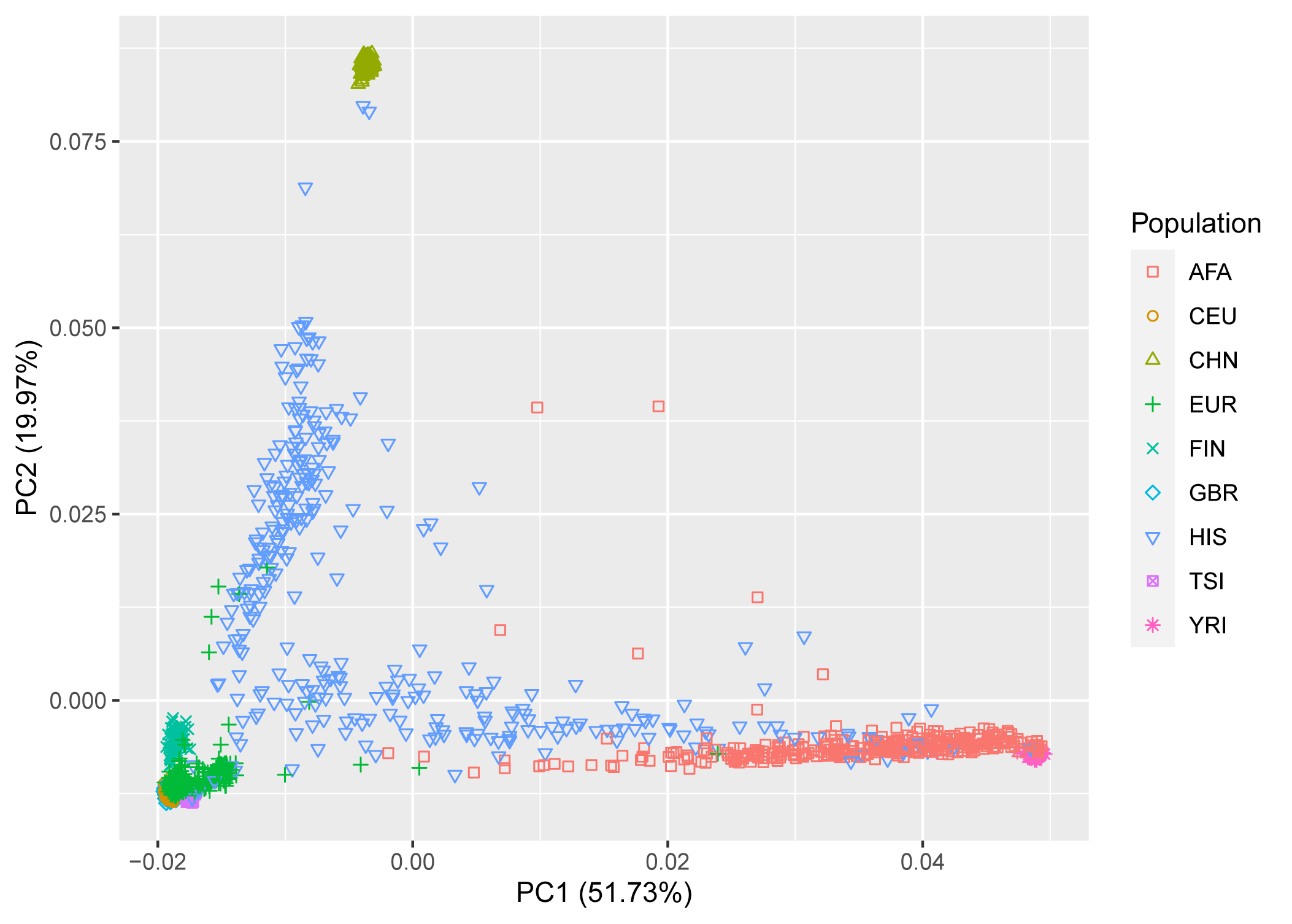

### Supplemental Figure 2

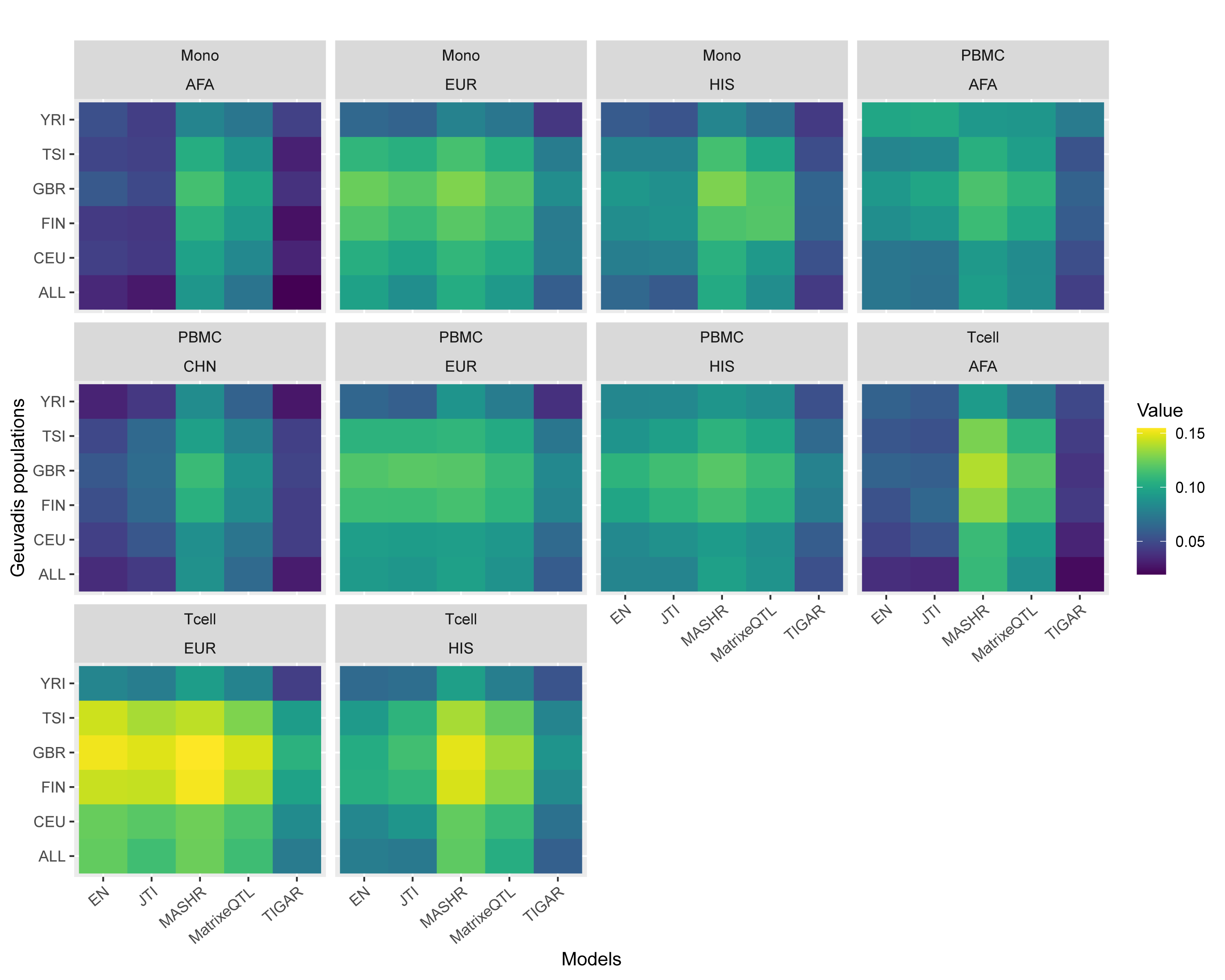

### Supplemental Figure 3

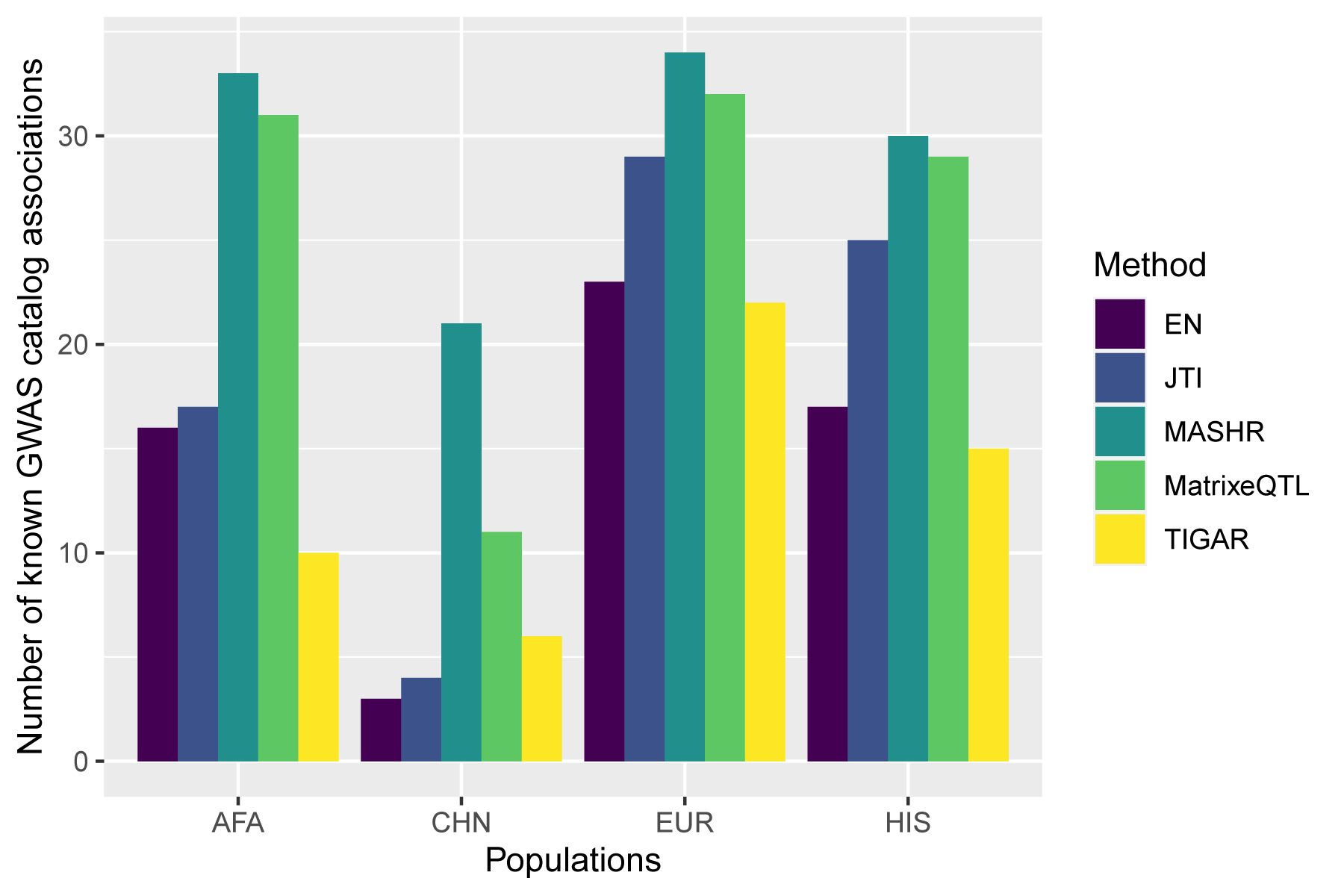
